# ALDH2 Knockout Induces Dysregulation of Arginine biosynthesis in SKBR-3 Breast Cancer Cells

**DOI:** 10.64898/2026.09.21.753265

**Authors:** Ming Zhao, Blake R Rushing, Zhikun Ma, Rachel A Coble, Amanda B Parris, Tationa Campbell, Sergey A Krupenko, Susan Sumner, Xiaohe Yang

**Author notes:** **Correspondence:** Xiaohe Yang Ph.D.

## Abstract

**Background/Objective:** Aldehyde dehydrogenase 2 (ALDH2), a mitochondrial enzyme, mitigates cellular stress by detoxifying reactive aldehydes produced during alcohol exposure and endogenous metabolic processes. ALDH2*2 polymorphism, which results in deficient catalytic activity, is prevalent in East Asian populations and is associated with an increased risk of cancer and other diseases. While the effects of ALDH2 deficiency on cellular stress responses are well documented, the specific metabolic dysregulations arising from this deficiency require further investigation.

**Methods:** An ALDH2 knockout subline of SKBR-3 cells was established using CRISPR/Cas9 strategy. The metabolomic profiles of control (SKBR-3/C) and ALDH2 knockout (SKBR-3/ADKO) cells were evaluated via ultra-high performance liquid chromatography-high resolution mass spectrometry (UHPLC-HRMS).

**Results:** ALDH2 knockout in SKBR-3 cells resulted in increased cellular stress, evidenced by elevated levels of 8-OHdG, reactive oxygen species (ROS), and malondialdehyde (MDA). Metabolomic analysis of the paired cell lines assigned a total of 5,033 signals, of which 281 were significantly different (fold change ≥ 2, p-value<0.05) between the two groups. Pathway analysis with MetaboAnalyst identified significant metabolic pathways associated with ALDH2 deficiency, particularly those involving arginine biosynthesis, and arginine and proline metabolism. ALDH2 deficiency reprogramed arginine metabolism toward polyamine pathways while suppressing the urea cycle. Network analysis further revealed alterations in amino acid and lipid metabolism in ALDH2-deficient cells.

**Conclusions:** This study provides a novel insight into the role of ALDH2 in cancer metabolism and may have broader implication for cancer biology.

## 1. Introduction

Aldehyde dehydrogenase 2 (ALDH2), a mitochondrial matrix-localized enzyme, serves as the primary catalyst for detoxifying both endogenous and exogenous acetaldehydes [1]. While ALDH2 is best known for its role in metabolizing acetaldehyde derived from alcohol consumption, it is also essential for the NAD^+^-dependent oxidation of endogenous aldehydes, such as 4-hydroxynonenal (4-HNE), a toxic byproduct of lipid peroxidation [2]. Functional ALDH2 is vital for maintaining cellular homeostasis by protecting cells from oxidative stress, DNA damage, and protein adduct formation. Beyond its detoxification functions, ALDH2 also plays a fundamental role in mitochondrial bioenergetics and redox balance, in part by modulating glutathione levels, an essential mechanism for regulating the cellular microenvironment [3,4].

ALDH2 deficiency, primarily caused by polymorphic variants with reduced enzymatic activities, represents a significant clinical issue. The most well-characterized variant is ALDH2*2 (rs671), which leads to markedly decreased enzyme activity, approximately 64% loss in heterozygotes and up to 95% in homozygotes. This enzyme variant, which has lysine at position 487 instead of glutamate, is prevalent in 30-45% of East Asian populations [5-7]. Extensive epidemiological evidence links ALDH2 deficiency to a broad spectrum of pathological conditions, including cardiovascular disease, neurodegenerative disorders and increased risk for certain types of cancers [8-10]. Studies using cell lines and ALDH2 knockout mouse models demonstrate that ALDH2 loss disrupts key physiological processes, particularly those involving oxidative stress, alcohol metabolism, and neurodegeneration. ALDH2 knockout mice exhibit significantly elevated acetaldehyde levels in blood and tissues, especially following alcohol feeding. This accumulation is associated with liver injury, cardiovascular dysfunction, and neurodegenerative changes, and has been shown to promote alcohol-induced liver cancer [11,12]. At a cellular level, ALDH2 loss was associated with heightened cellular stress responses, increased ROS production, DNA damage, and altered signaling pathways, including MAPK, JNK, and p53 activation [13,14]. Understanding the molecular mechanisms underlying ALDH2 deficiency is critical for addressing its significant health impacts.

Although alcohol consumption is a known risk factor for breast cancer, the role of ALDH2 deficiency in mammary tumorigenesis remains unclear. Epidemiological studies have produced mixed results on this issue. The ALDH2 rs671 polymorphism, which significantly reduces enzyme activity, has been linked to increased risk for several alcohol-related cancers [15]. A pooled analysis of 14 case-control studies in Asian women found that the homozygous rs671 genotype was associated with higher risk for breast cancer, particularly hormone receptor-positive subtypes [16]. Another ALDH2 SNP, rs10744777, has been shown to modify breast cancer risk in carriers of the low-penetrance BRCA2 variant, suggesting that impaired ALDH2 function, and the resultant aldehyde accumulation, may interact with altered BRCA2 activity to increase susceptibility [17]. However, other studies have reported no significant association between ALDH2 polymorphisms and overall breast cancer risk, highlighting the complexity of this relationship and the influence of potential confounding factors [18]. These inconsistencies may be partly due to variations of participant’s conditions across different studies. Of note, there is a lack of preclinical models to directly investigate the role of ALDH2 deficiency in breast cancer independent of alcohol exposure. To address this gap, we developed ALDH2-specific knockout models to explore its role in breast cancer pathogenesis.

Specifically, we compared isogenic HER2-positive SKBR-3 breast cancer cell lines with and without ALDH2 expression. These cells were selected for their high endogenous ALDH2 expression and clinical relevance to aggressive breast cancer subtypes. Using CRISPR/Cas9-mediated gene editing, we established paired control and ALDH2-knockout cell lines and integrated functional assessments of oxidative stress with high-resolution untargeted metabolomics to characterize the global metabolic signature of ALDH2 deficiency, to identify dysregulated pathways driving phenotypic changes, and to uncover novel metabolic vulnerabilities of breast cancer. Our study provides the evidence that ALDH2 loss is associated with remodeling of arginine, redox, and lipid-linked metabolism. These findings establish ALDH2 as a critical metabolic regulator and suggest targetable nodes for therapeutic intervention in ALDH2-compromised tumors.

## 2. Materials and Methods

### 2.1. Reagents

SKBR-3 cells were purchased from the American Type Culture Collection (ATCC) (Manassas, VA, USA). All solvents for UHPLC-HRMS analysis (Optima-grade water and methanol with 0.1% formic acid) were purchased from Fisher Scientific (Waltham, MA, USA). Dulbecco’s Modified Eagle Medium/Nutrient Mixture F-12 (DMEM/F-12) was purchased from Gibco (Grand Island, NY, USA). Fetal bovine serum (FBS) was purchased from GeminiBio (West Sacramento, CA). Bicinchoninic acid kit (BCA) assays were purchased from Thermo Scientific (Madison, WI, USA). Laemmli sample buffer was purchased from Bio-Rad (Hercules, CA). Primary antibody ALDH2 (Cat:MA5-17029) was from Invitrogen. ALDH1A1 antibody (Cat: 15910-1-AP) was from Proteintech (Rosemont, IL). β-Actin antibody (Cat: sc-47778) were from Santa Cruz Biotechnology (Dallas, USA). horseradish peroxidase (HRP)-conjugated Mouse/Rabbit secondary antibodies were from Cell Signaling Technology (Danvers, MA).

### 2.2. Cell culture

SKBR-3 cells were maintained in DMEM/F-12 medium supplemented with 10% FBS, 100 U/mL penicillin, and 100 μg/mL streptomycin. Cells were grown in a 37 °C, 5% CO2, humidity-controlled environment. To prepare cells for metabolomics analysis, 4×10^6^ SKBR-3 cells were plated into 100 mm tissue culture dishes and allowed to adhere overnight, achieving approximately 80% confluency.

### 2.3. Generation of ALDH2-knockout cell line

Lentiviral ALDH2-knockout vector HCP349485-LvSG01-3 and empty vector were all from Genecopoeia (Rockville, MD), and were used to generate the paired control and ALDH2 knockout cell lines. For lentivirus packaging, 293T cells were transfected with a lentivirus packing kit (Genecopoeia, Rockville, USA). After infection of 2 days, the cells were screened with 2 μg/mL puromycin for one weeks. The expected ALDH2 knockout clone was selected and applied for the study.

### 2.4. ROS measurement

CM-H2DCFDA detection kit was used to determinate ROS levels in cells according to manufacturer’s instruction. Briefly, the cells in log phase were harvested with trypsinization and suspended in PBS, and then incubated with 10 μM DCF-DA at 37 °C for 30 min. the cells were collected with centrifugation, washed with PBS three times. Green fluorescence data were measured using a Guava EasyCyte 8 flow cytometer.

### 2.5. MDA assay

The levels of malondialdehyde (MDA) were determined using lipid peroxidation (MDA) assay kit (ab118970, Abcam, Waltham, MA). Briefly, 2×10^6^ cells in each group was collected and then homogenized in lysis solution. After centrifugation at 13,000×*g* for 10 min. The supernatant was collected. The mixture of Developer/TBA reagent with sample or standard were incubated at 95°C for 10 min. Then, cool room temperature for 10 min. 200 µl of the reaction mix were transferred into 96-well microplate for analysis at OD 532 nm.

### 2.6. Immunofluorescence (IF)

The cells were fixed in 4% paraformaldehyde for 15 min at room temperature. After washing with PBS, the cells were permeabilized with 0.1% Triton X-100 for 20 min and continually blocked with 3% BSA (in PBST) for 30 min. Cells were then incubated with primary antibodies diluted in 3% BSA overnight at 4 °C. After washing three times, Alexa Fluor 594-labeled secondary antibody (Thermo Fisher Scientific) diluted in 3% BSA solution were added and incubated for 1 h at room temperature in the dark. After washing three times, the cells were mounted in Antifade Mounting Medium containing DAPI (VecorLabs). Images were acquired using a Nikon Microscope.

### 2.7. Western blot

The cells were collected with trypsinization, and lysed in Laemmli sample buffer (Bio-Rad, Hercules, CA), followed by protein concentration determination with a BCA assay kit. Thirty (30) μg of protein lysate from each sample was loaded for separation by SDS-PAGE electrophoresis. Separated proteins were transferred to polyvinylidene difluoride (PVDF) membranes. The membrane was blocked with 5% non-fat milk in TBST at room temperature for 1 h, and then incubated with specific primary antibodies at appropriate dilutions at 4°C overnight. After washing with TBST for 3 times, the membranes were incubated with HRP-labeled secondary antibody (1:2000) for 1 h. The specific proteins were detected with chemiluminescence using the Super Signal West Dura Detection System (Thermo Fisher Scientific, Waltham, USA) and visualized by an Azure Biosystems Imager (Dublin, USA).

### 2.8. Quantitative real-time PCR (*q*-PCR)

Total RNA was isolated with TRIzol Reagent. iScript cDNA Synthesis kit (Bio-Rad) was used for reverse transcription. SYBR green-based q-PCR assays were performed in triplicate using a Bio-Rad CFX96 PCR machine. The primers for ALDH2, ALDH1A1and GAPDH in the *q*-PCR assay are: ALDH2, F: 5’-atcatgcaagcttcctccct-3’, R:5’-actcttaccctcagccaacc-3’; ALDH1A1, F: 5’-gttgtcaaaccagcagagca-3’, R:5’-cttttcccggcagcttcttt-3’; GAPDH, F: 5’-ctgacttcaacagcgacacc-3’, R: 5’-gtggtccaggggtcttactc-3’. The comparative-Ct method (ΔΔCt method) was used to calculate relative mRNA levels of individual samples.

### 2.9. Metabolite Extraction

At endpoint, dishes were placed on ice. Then, the cells were washed twice with 10 ml of ice-cold PBS and quenched with 1 mL of ice-cold acetonitrile. After 750 µL of ice-cold water was added to dishes, the cells were dislodged by scraping, and cell suspensions were transferred to 15 mL conical tubes. The remaining cell suspension was collected with the above same procedures. The two batches were combined to yield a total of approximately 3.5 mL of extract per sample. After vortexing at 6,000× *g* for 10 min, and then clarified by centrifugation at 16,000× *g* for 10 min at 4 °C. Supernatants were dried in speed-vac and reconstituted in 95:5 water: methanol proportionally to each sample’s protein concentration determined by BCA assay. A quality control study pool (QCSP) was created by mixing 10 μl of each sample.

### 2.10. Ultra High-Performance Liquid Chromatography-High Resolution Mass Spectrometry (UHPLC-HRMS) Analysis

Metabolomics data were acquired with a Vanquish UHPLC system coupled to a Q Exactive™ HF-X Hybrid Quadrupole-Orbitrap Mass Spectrometer (Thermo Fisher Scientific) using previously described methods [19]. Cell samples were randomized and injected onto the UHPLC-HRMS platform with QCSP every 8 samples in each group. Separation of metabolites was performed using a HSS T3 C18 column (2.1 × 100 mm, 1.7 µm, Waters Corporation) at 50 °C with binary mobile phase of water (A) and methanol (B), each containing 0.1% formic acid (v/v). The UHPLC linear gradient started from 2% B, and increased to 100% B in 16 min, then held for 4 min, with a flow rate of 400 µL/min. The untargeted data were acquired from 70 to 1050 m/z using data-dependent acquisition mode. Peak picking, alignment, and normalization was performed using Progenesis QI (version 2.1, Waters Corporation, Milford, MA, USA). QCSP and method blanks were analyzed after every six study samples to evaluate instrument stability and performance throughout data acquisition. Background removal was performed by filtering out peaks with a higher average abundance in the blank injections as compared to the QCSP injections. Peaks were then normalized in Progenesis QI using the “normalize to all” feature. Principal component analysis (PCA) and orthogonal partial least squares-discriminant analysis (OPLS-DA) were performed using SIMCA 16 (Umetrics, Umeå, Sweden). Data quality was assessed by visualizing the clustering and centering of QCSP injections with the study samples in PCA plots. Strong OPLS-DA models were defined as having R2X, R2Y, and Q2 > 0.5.

### 2.11. Metabolite Identification/Annotation and Metabolite Analysis

Identification and annotation of peaks to metabolites was performed by matching to an in-house reference standard RT, Mass, MS/MS library of over 2400 compounds run on the UHPLC-HRMS platform, or to public databases (NIST, METLIN, HMDB). Metabolite assignments were based on matches of peaks to exact mass (MS, <5 ppm), MS/MS fragmentation pattern (similarity score > 30%), isotopic ion pattern (similarity score > 90%), or retention time (RT, for in-house library standards only, ±0.5 min). OL1 refers to an in-house library match by MS, MS/MS, and RT; OL2a refers to an in-house library match by MS and RT; OL2b refers to an in-house library match by MS and MS/MS; PDa refers to a public database match by MS and experimental MS/MS (NIST or METLIN); PDb refers to a public database match by MS and theoretical MS/MS (HMDB); PDc refers to a public database match by MS and isotopic similarity; PDd refers to a public database match by MS only. Metabolites matched at the OL1 and OL2a level were used for analytes conducted in MetaboAnalyst 6.0. The names given for each match are based on the names of the reference standards run on our UHPLC-HRMS platform or the names provided in public databases. This method does not necessarily differentiate between some isomeric forms such as D and L enantiomers. Network visualization was performed using the Metscape plug-in for Cytoscape as previously[19,20]. Fold changes were calculated using median peak abundance values and p-values were calculated using Student’s t-test. Pathway analysis was performed using the Enrichment Analysis module of MetaboAnalyst 6.0. Only metabolites matched to the in-house library at a level of OL1 and OL2a with an VIP>1 in knockout vs control samples were used for the analysis. Pathway analysis was performed separately for increased or decreased metabolites in the knockout vs control comparison.

### 2.12. Statistical analysis

Quantitative results were statistically analyzed using GraphPad Prism 9 software. Comparisons between two groups in single-factor experiments were performed using a two-tailed Student’s t-test. ** p < 0.01.

## 3. Results

### 3.1. Specific knockout of ALDH2 in SKBR-3 cells establishes a model for functional and metabolic analysis

To investigate the effect of ALDH2 deficiency on cellular function and the associated metabolic disturbance in breast cancer cells, we compared paired cell lines with and without ALDH2 expression. We first examined the expression levels of ALDH2 in a panel of breast cancer cell lines, including T47D, MCF-7, SKBR-3, BT-474 and MDA-MB-231 cell lines, for the selection of the cell line to be used for ALDH2 knockout. As shown in Fig.1A, among five breast cancer cell lines, ALDH2 protein levels are higher in SKBR-3, BT-474 and MDA-MB-231 cells, as compared to T47D and MCF-7 cells. To study the effect of ALDH2 deficiency on cellular functions and metabolic disturbance, we generated stable control and ALDH2 specific knockout sublines based on SKBR-3 cell line, which has relatively high ALDH2, using lentivirus mediated CRISPR/Cas9 technology. As shown in Fig. 1B, ALDH2 knockout was efficient in the ALDH2 knockout subline (SKBR-3/ADKO), as compared to the control (SKBR-3/C). To demonstrate the knockout was ALDH2 specific, we examined the expression of ALDH1A1, another major member of the ALDH family. Results from both protein and mRNA levels indicated that ALDH1A1 expression was not affected in the SKBR-3/ADKO cells. These results indicate we established ALDH2 knockout sublines ready for subsequent functional and metabolic studies.

**Figure 1.**
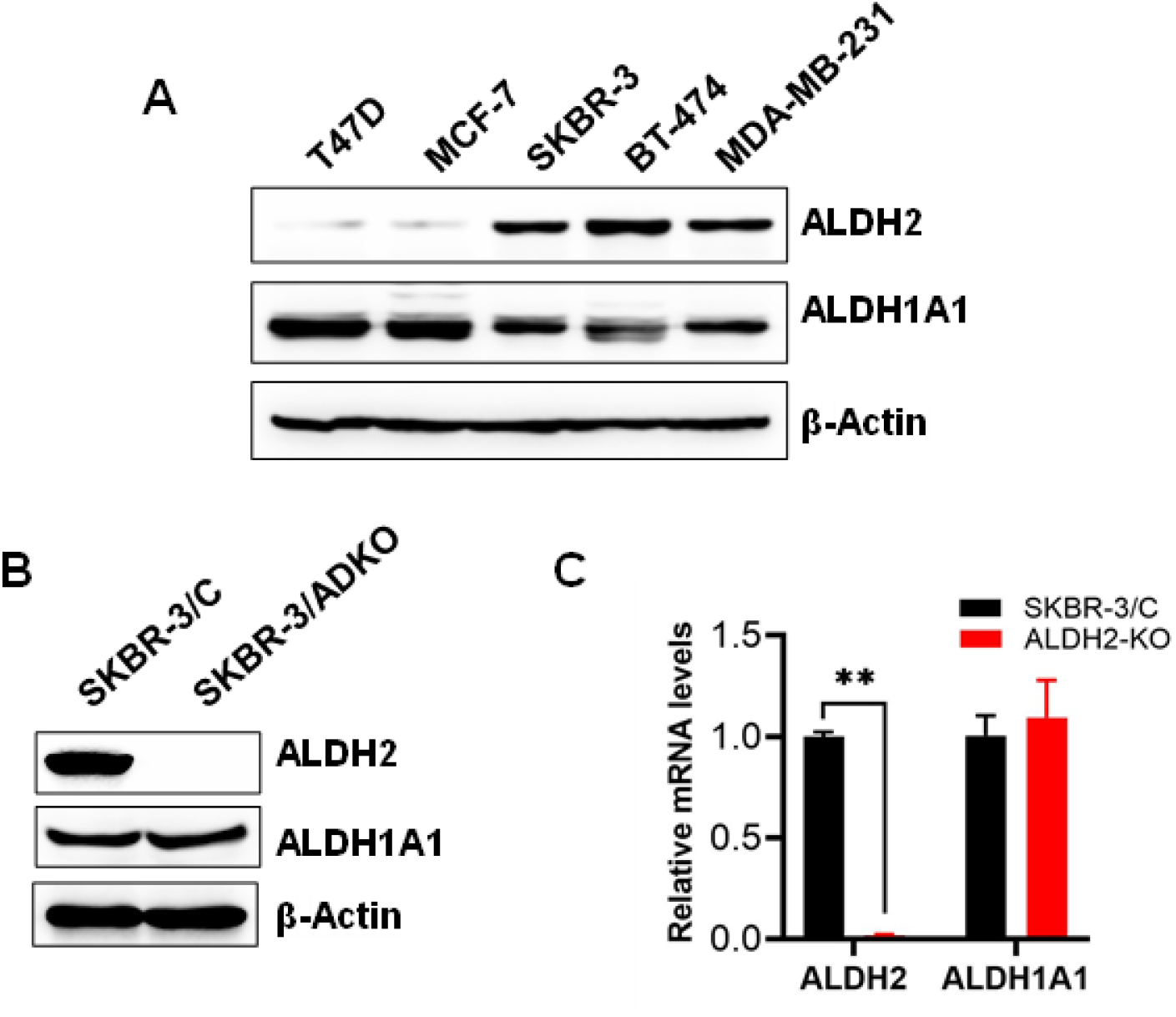
ALDH2 was specifically knocked out in SKBR-3 cells. A. Protein levels of ALDH2 and ALDH1A1 in five breast cancer cell lines (T47D, MCF-7, SKBR-3, BT-474, MDA-MB-231). β-Actin serves as the internal control. B. Protein levels of ALDH2 and ALDH1A1 in control (SKBR-3/C) and ALDH2 knockout (SKBR-3/ADKO) cells were de-termined by Western blotting. C. mRNA levels of ALDH2 and ALDH1A1 in SKBR-3/C and SKBR-3/ADKO cells were measured by q-PCR. **p<0.01.

### 3.2. ALDH2 knockout induces cellular stress in SKBR-3 Cells

To assess the functional impact of ALDH2 knockout in SKBR-3 cells, we evaluated its effects on cellular oxidative stress, a hallmark of ALDH2 dysregulation, using multiple approaches. Immunofluorescence staining revealed significantly increased levels of 8-OHdG, a key marker of oxidative DNA damage, in ALDH2 knockout cells compared to control cells (Fig. 2A & B). Similarly, the malondialdehyde (MDA) content, an indicator of lipid peroxidation, was markedly higher in ALDH2 knockout cells (Fig. 2C). We further quantified intracellular ROS levels using a CM-H2DCFDA-based assay and flow cytometry. Consistent with other findings, ALDH2 knockout cells displayed significantly elevated ROS levels compared to controls (Fig. 2D). These results highlight the critical role of ALDH2 in maintaining cellular homeostasis and underscore the functional impact of the enzyme loss. These findings also indicate that we generated a suitable model to address the functional role of ALDH2 in regulating cellular metabolism.

**Figure 2.**
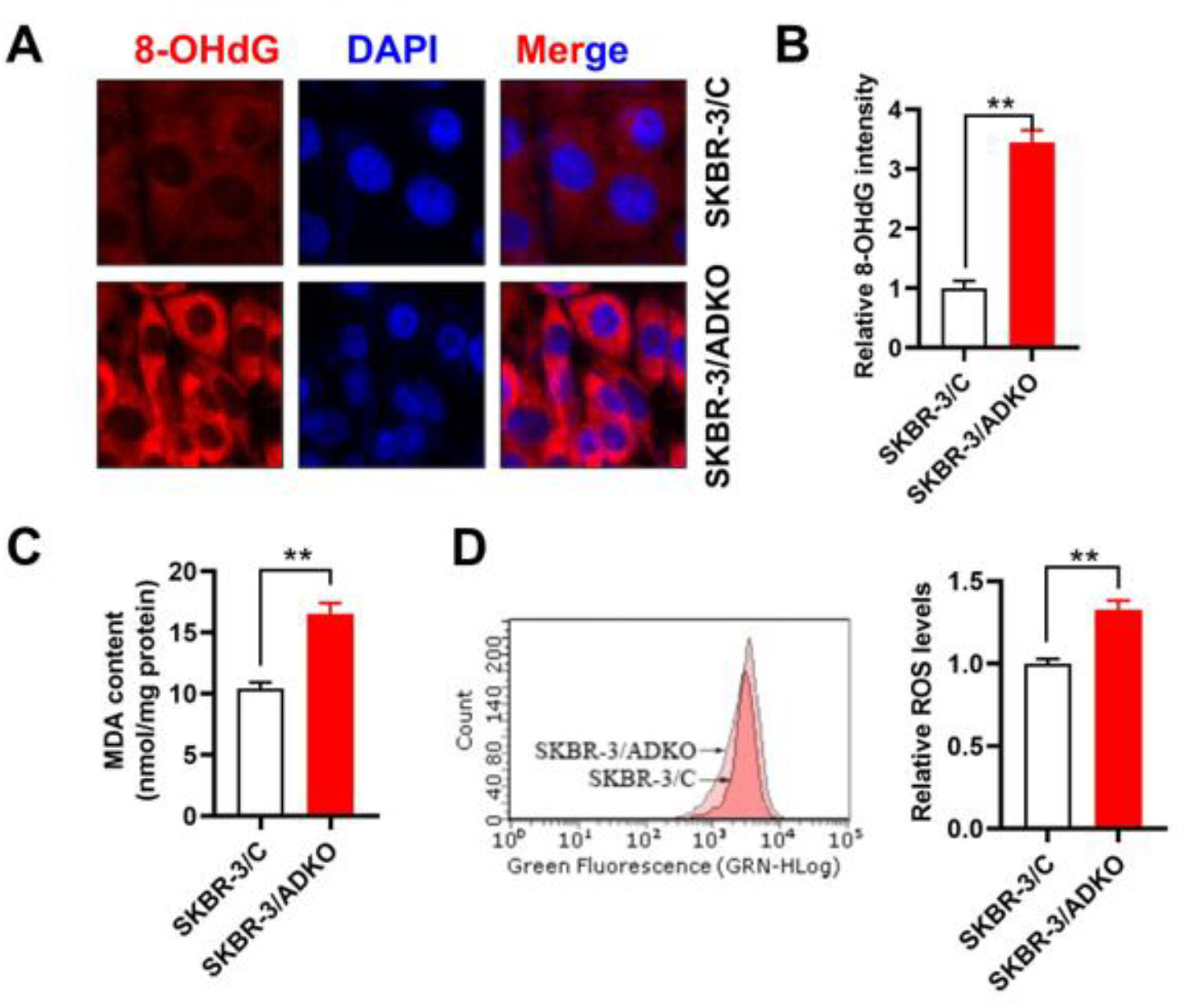
ALDH2 knockout induces oxidative stress in SKBR-3 cells. A & B. Representative images of 8-OHdG immunofluorescence staining in SKBR-3/C and SKBR-3/ADKO cells. 8-OHdG and nuclei are labeled with red and blue, respectively. The bar graph shows quan-titative analysis of fluorescence intensity, **p<0.01. C. MDA levels in paired cell lines are measured using a colorimetric kit. data are presented as means ± SEM from three inde-pendent replicates, **p<0.01. D. ROS levels in SKBR-3/C and SKBR-3/ADKO cells are de-termined by CM-H2DCFDA staining and flow cytometry. Representative overlap histograms are shown. The bar graph represents mean fluorescence intensity (means ± SEM, n=3, **p<0.01.

### 3.3. ALDH2 knockout induces distinctive metabolomic profile in SKBR-3 cells

To determine the impact of ALDH2 knockout on the metabotype of SKBR-3 cells, we conducted an untargeted metabolomics analysis using the LC-MS/MS platform. A total of 5033 peaks remained after preprocessing and 3869 were identified and annotated. Among these, 266 metabolites were assigned with OL1 and OL2a confidence and an additional 289 metabolites matched as OL2b and PDA (Table S1). The principal component analysis (PCA) score plot revealed a clear metabolite separation between the control and knockout groups (Fig.3A), indicating that there were distinct metabolic signatures between the two cell lines. Supervised orthogonal partial least squares-discriminant analysis (OPLS-DA) was used as an additional method to evaluate such differences. The prediction ability (Q2) of 0.865 further demonstrated that ALDH2 knockout resulted in the change of metabolic profile (Fig.3B). Additionally, heatmap analysis illustrates distinctive patterns of metabolomic profiles between the paired sublines (Fig. 3C). These results demonstrate that ALDH2 knockout has a profound impact on the metabolism of SKBR-3 cells.

**Figure 3.**
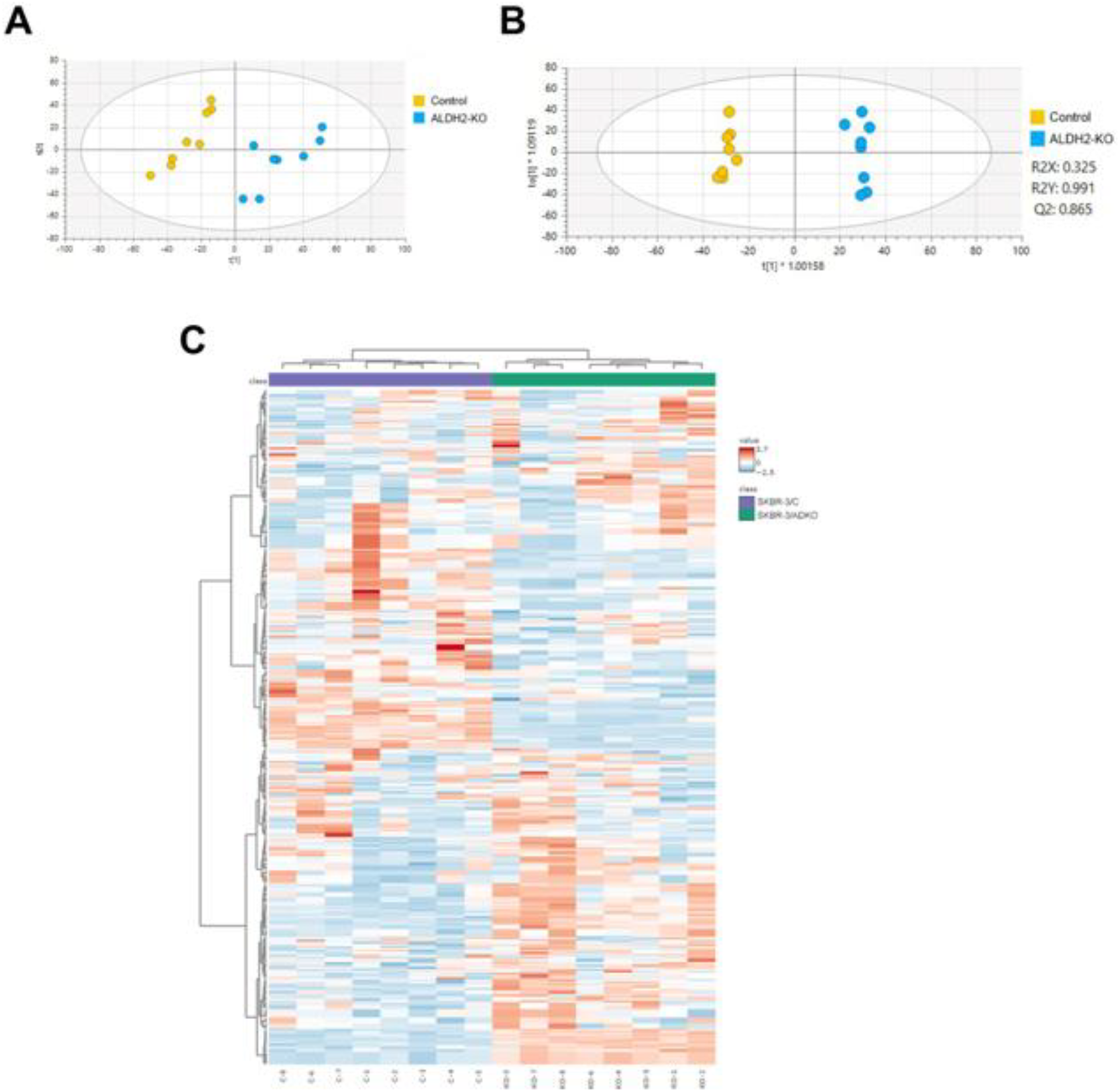
Multivariate analysis of metabolomic profiles in SKBR-3/C and SKBR-3/ADKO cells. A. Principal Component Analysis (PCA) of metabolomic profiles. B. Orthogonal partial least squares discriminant analysis (OPLS-DA) of metabolomic data, with model parameters: R^2^X = 0.325, R^2^Y = 0.991, and Q^2^ = 0.865. C. Heatmap of the top 100 differentially expressed metabolites between SKBR-3/C and SKBR-3/ADKO cells, generated using MetaboAnalyst 6.0. The data were derived from 5033 detected peaks (Supplementary Table S2, n = 8).

### 3.4. Loss of ALDH2 drives metabolic reprogramming associated with altered fatty acid oxidation and amino acid metabolism

To identify the top differentially regulated metabolites with a visualized presentation, we analyzed the metabolomics data from the control and ALDH2 knockout groups using a volcano plot. This plot highlights the most significant metabolites based on both fold-change and statistical significance (fold change ≥ 2, p < 0.05). As shown in Fig. 4, the plot provides a clear overview of significantly regulated metabolites induced by ALDH2 knockout, with the top 20 altered metabolites labeled. Among these, several carnitine derivatives, such as myristoyl-L-carnitine, octadecanoylcarnitine, 3-hydroxyhexadecanoylcarnitine, hexadecanoylcarnitine, dodecanoylcarnitine, and linoleyl carnitine, were markedly downregulated in ALDH2 knockout cells. Other significantly downregulated metabolites included L-citrulline, thymine, histidine, ureidosuccinic acid, iminodiacetic acid, spermine, and galactose. Conversely, significantly upregulated metabolites included glycylvaline, methylthioadenosine, glycerophosphocholine, lysyl-leucine, 7alpha-hydroxy-3-oxo-4-cholestenoic acid, 3-hydroxyglutarate, and docosahexaenoic acid. A more comprehensive list of metabolites identified through volcano plot analysis is provided in supplementary files (Table S2).

**Figure 4.**
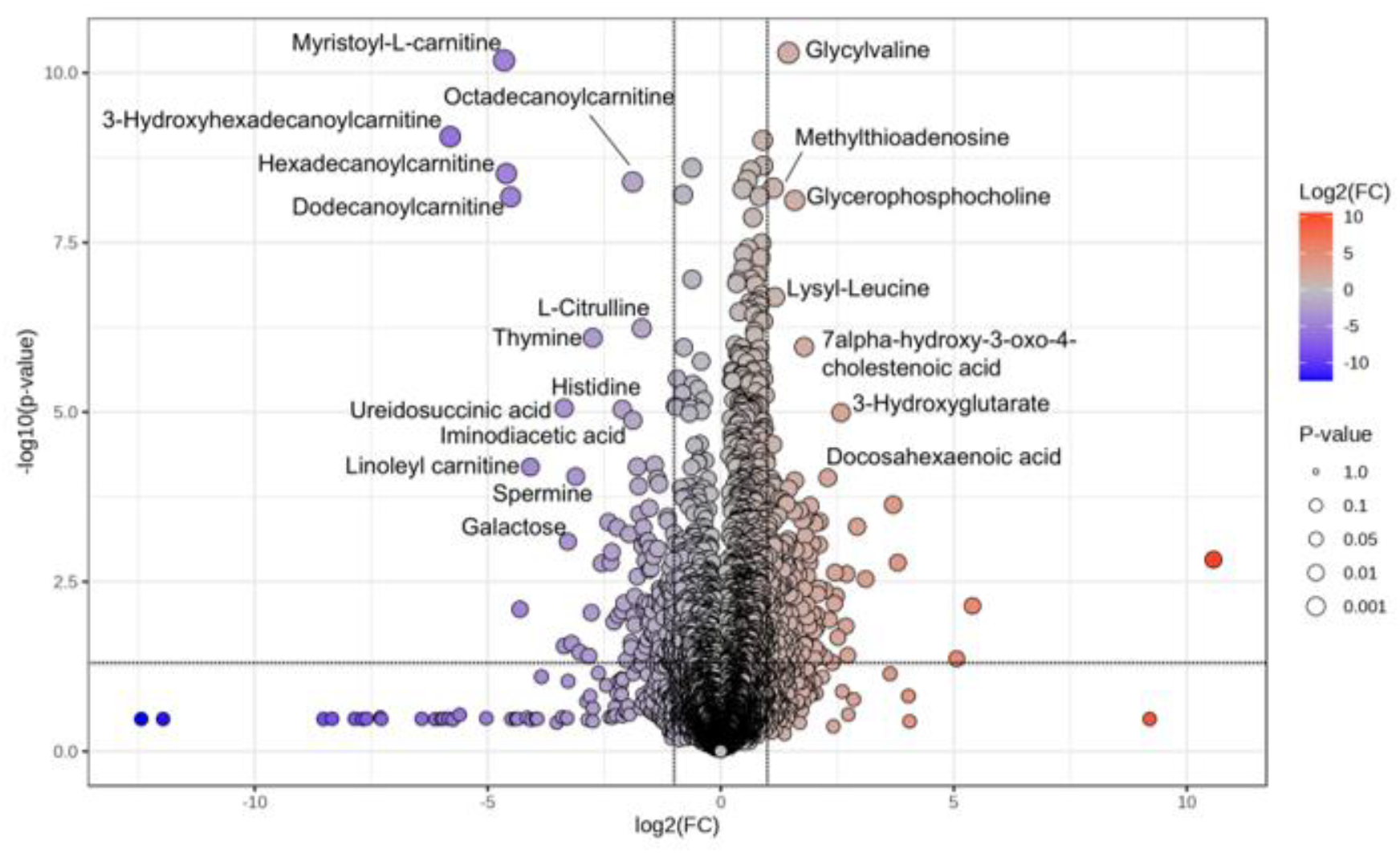
Volcano plot of differentially expressed metabolites between SKBR-3/C and SKBR-3/ADKO cells. The volcano plot displays metabolite differences between groups, with log-transformed adjusted p-values on the y-axis and fold changes on the x-axis. Upregulated metabolites are highlighted in red, downregulated metabolites blue (fold change≥2, p<0.05), and non-significant metabolites in grey. The top 20 significant metabolites are labeled.

### 3.5. ALDH2 deficiency reprograms amino acid and redox metabolism, highlighting arginine and glutathione pathways

To understand the effect of ALDH2 deficiency on specific metabolic pathways, we analyzed the metabolite profiles with MetaboAnalyst 6.0. Based on variable importance in projection (VIP) scores, metabolites with VIP>1 were subjected to pathway enrichment analysis (Table S3). The top enriched pathways included arginine biosynthesis, arginine and proline metabolism, alanine, aspartate and glutamate metabolism, glutathione metabolism, and phenylalanine, tyrosine and tryptophan biosynthesis (Fig. 5A, Table S4). Among these, arginine biosynthesis and arginine-related pathways showed the strongest statistical significance, indicating a prominent impact on nitrogen handling and urea cycle-linked metabolism. These pathways are biochemically interconnected and central to cellular metabolic homeostasis. Arginine and proline metabolism are closely linked to nitric oxide production, polyamine synthesis, and mitochondrial function, while alanine, aspartate, and glutamate metabolism play key roles in transamination reactions and anaplerotic input into the tricarboxylic acid (TCA) cycle. The enrichment of glutathione metabolism suggests altered redox balance, consistent with the role of ALDH2 in detoxifying reactive aldehydes and limiting oxidative stress. Additionally, perturbation of aromatic amino acid biosynthesis pathways may reflect broader changes in biosynthetic and signaling processes. To further examine metabolite patterns within these pathways, a heatmap analysis was performed. As shown in Fig. 5B, metabolites within these enriched pathways display distinct clustering and differential abundance between control and ALDH2 knockout cells, indicating coordinated metabolic reprogramming. Collectively, these results demonstrate that ALDH2 deficiency reshapes interconnected amino acid and redox metabolic networks.

**Figure 5.**
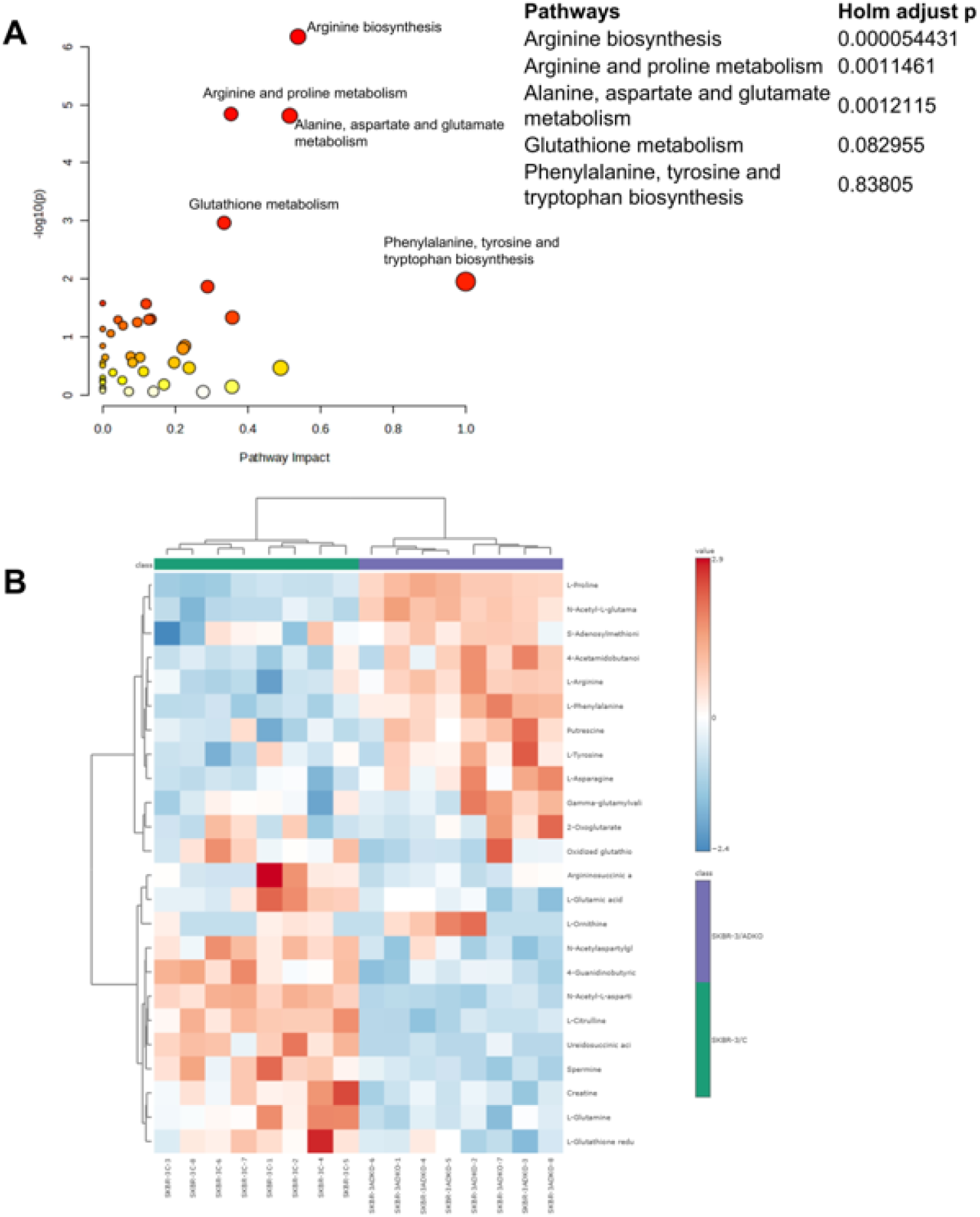
ALDH2 knockout induces differential regulation of metabolic pathways in SKBR-3 cells. A. Pathway enrichment analysis comparing SKBR-3/C and SKBR-3/ADKO cells. The top five pathways, ranked by Holm-adjusted p-values, are shown. Circle color in-tensity and size indicate lower p-values and higher numbers of significantly altered metabolites relative to pathway size, respectively. B. Heatmap of differentially expressed metabolites involved in the top five enriched pathways.

### 3.6. ALDH2 deficiency redirects arginine metabolism toward polyamine pathways with suppression of the urea cycle

To quantitatively assess metabolite alterations within the top enriched pathways, we performed supervised analysis of key metabolites in each pathway and visualized their fold changes based on metabolite intensities (Fig. 6A, Table S5). Distinct and pathway-specific patterns emerged, indicating coordinated metabolic reprogramming in ALDH2-deficient cells. Within the arginine biosynthesis pathway, key intermediates of the urea cycle, including L-citrulline and argininosuccinic acid, were markedly decreased, whereas upstream and branch-point metabolites such as L-ornithine, N-acetyl-glutamate, and L-arginine were significantly increased. This pattern suggests a disruption of canonical urea cycle flux. In parallel, arginine and proline metabolism showed a pronounced reduction in spermine, consistent with earlier observations (Fig. 4), alongside an increase in putrescine, indicating an imbalance in polyamine metabolism. More broadly, metabolites within the alanine, aspartate, and glutamate metabolism pathway were predominantly downregulated, suggesting reduced anaplerotic input into the TCA cycle and altered nitrogen redistribution. In the glutathione pathway, opposing trends in polyamine-related metabolites (e.g., decreased spermine and increased putrescine) further support perturbation of redox-associated metabolic networks. In contrast, aromatic amino acids (L-tyrosine and L-phenylalanine) showed modest but consistent downregulation, indicating a more limited effect on these biosynthetic pathways. Given that arginine biosynthesis was the most significantly affected pathway (Fig. 5A), we further mapped metabolite changes onto the KEGG arginine biosynthesis pathway (Fig. 6B) to assess pathway flux. Upstream metabolites such as 2-oxoglutarate (1.13-fold) and L-glutamic acid (0.86-fold) exhibited relatively minor changes, whereas L-ornithine (1.70-fold), N-acetyl-glutamate (1.29-fold), and L-arginine (1.30-fold) accumulated substantially. Notably, downstream metabolites in the polyamine branch, including putrescine (1.80-fold) and N4-acetylaminobutanoic acid (1.24-fold), were increased, while spermine was markedly decreased (0.29-fold). In contrast, intermediates of the urea cycle, including L-citrulline and argininosuccinic acid, were significantly reduced (0.31-fold and 0.47-fold, respectively), suggesting impaired flux through the argininosuccinate synthase/lyase steps. Collectively, these findings indicate a metabolic shift in which accumulated L-ornithine is preferentially diverted away from the urea cycle toward polyamine synthesis and related acetylated derivatives. This rewiring of arginine metabolism highlights a potential mechanism by which ALDH2 deficiency alters nitrogen handling, redox balance, and proliferative metabolic pathways.

**Figure 6.**
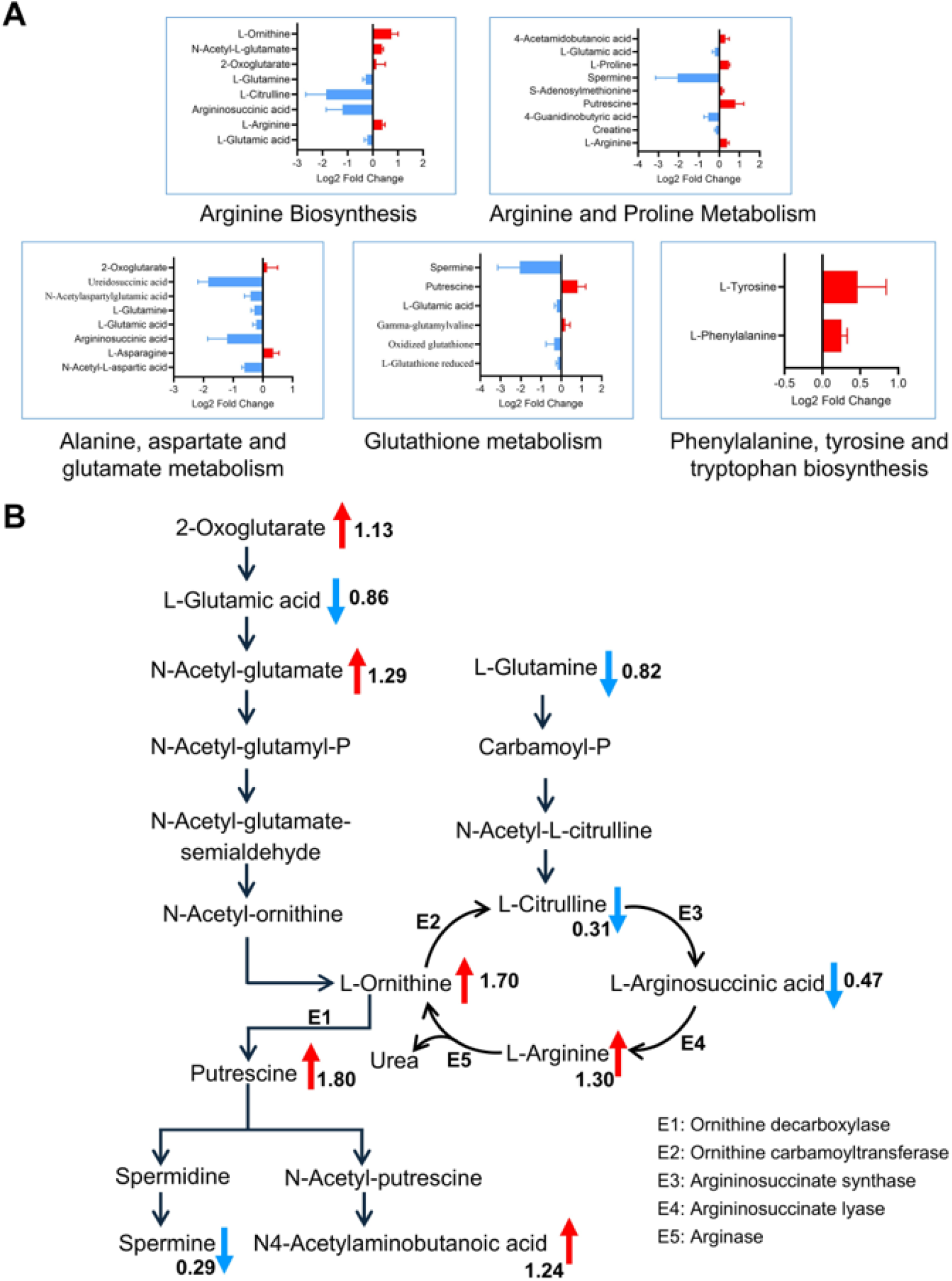
Supervised analysis of key metabolites in enriched pathways. A. Changes in key metabolites within enriched pathways induced by ALDH2 knockout. Metabolites were extracted and plotted to show log2 fold changes based on normalized intensities. B. Integration of altered metabolites in the arginine and proline metabolism pathways. The pathway diagram was adapted from the KEGG database.

### 3.7. Network analysis reveals integrated reprogramming of amino acid and lipid metabolism in ALDH2-deficient cells

To further investigate the relationships among altered metabolites and to integrate pathway-level findings, we performed metabolic network analysis using Metscape based on significantly differentiated metabolites (VIP>1). The resulting network (Fig.7) revealed interconnected clusters of metabolic pathways, highlighting coordinated remodeling of cellular metabolism in ALDH2-deficient cells. Consistent with pathway enrichment analysis (Fig.5), amino acid metabolism formed a central hub within the network. Prominent clusters included glycine, serine, alanine, and threonine metabolism, as well as arginine, proline, glutamate, aspartate, and asparagine metabolism, reflecting extensive perturbation of nitrogen metabolism and amino acid interconversion pathways. Additional subnetworks involving histidine, tyrosine, and tryptophan metabolism further indicate widespread remodeling of amino acid-derived biosynthetic and signaling processes. In parallel, multiple lipid-related pathways were enriched and interconnected within the network, including glycerophospholipid metabolism, arachidonic acid metabolism, and cholesterol biosynthesis (via squalene). The enrichment of nicotinate and nicotinamide metabolism suggests potential alterations in NAD^+^-dependent redox processes, which is consistent with the known role of ALDH2 in aldehyde detoxification and redox homeostasis. Notably, the network structure highlights extensive crosstalk between amino acid and lipid metabolic pathways, suggesting that ALDH2 deficiency induces coordinated metabolic reprogramming rather than isolated pathway changes. These findings extend the pathway-level observations by placing the altered metabolites into a systems-level context, supporting a model in which disruptions in nitrogen metabolism, redox balance, and lipid biosynthesis are functionally interconnected.

**Figure 7.**
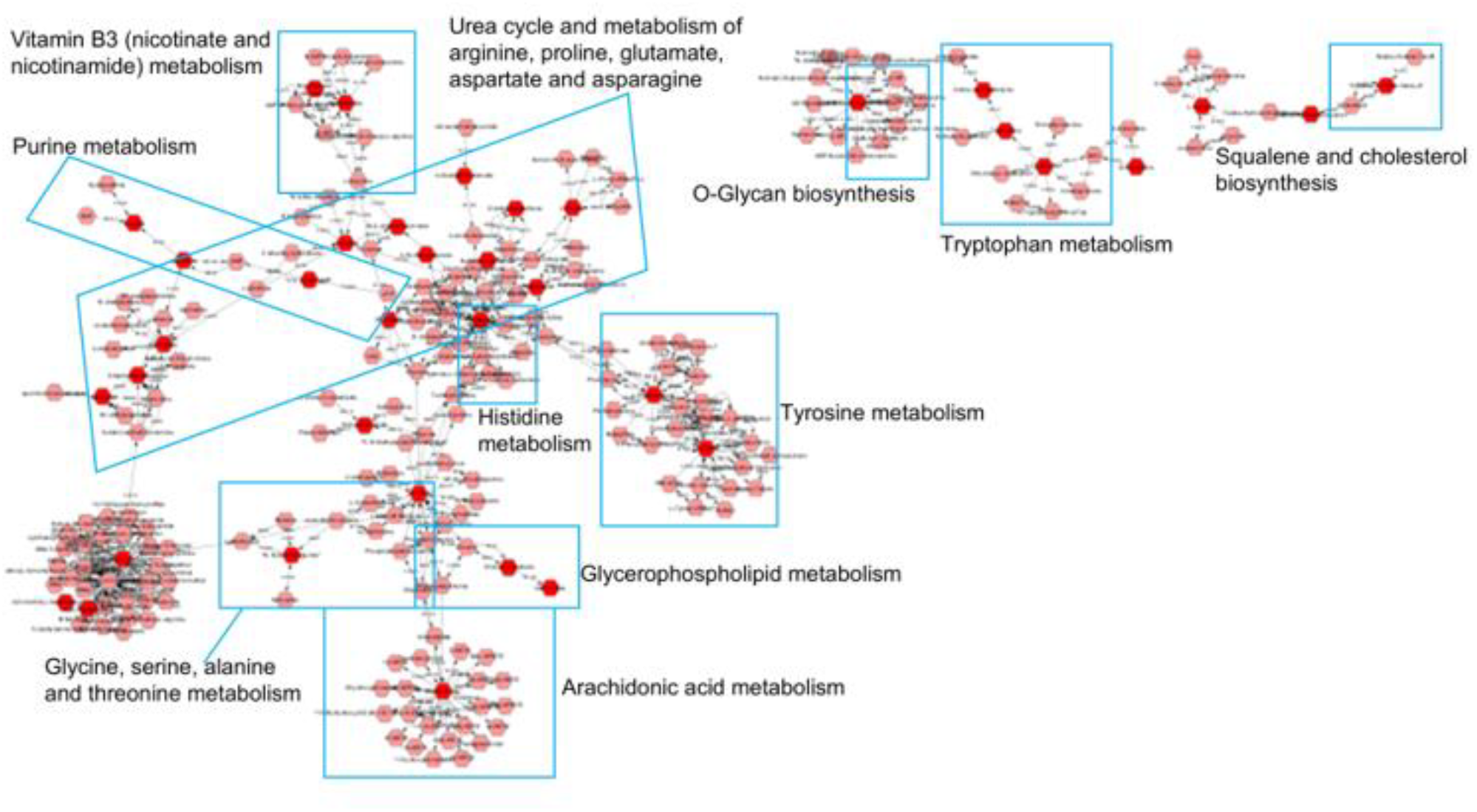
Network analysis of metabolic pathways affected by aldh2 knockout. The ALDH2-regulated metabolic network was constructed by comparing SKBR-3/C and SKBR-3/ADKO cells. Differential metabolites at OL1 and OL2a levels were analyzed using Metscape. Metabolites are organized into pathways based on the KEGG human database. Hexagons represent metabolites, and edges indicate interactions. Light red nodes denote metabolites without significant changes, while dark red nodes represent significantly altered metabolites.

## 4. Discussion

This study aimed to evaluate the impact of ALDH2 deficiency on cellular stress and associated metabolomic alterations in breast cancer cells. Using isogenic SKBR3 sublines generated in this study, we demonstrated that loss of ALDH2 is sufficient to induce a pronounced cellular stress phenotype accompanied by coordinated metabolic rewiring. Specifically, ALDH2 knockout increases intracellular ROS levels and elevates malondialdehyde (MDA), a marker of lipid peroxidation, as well as 8-OHdG, an indicator of oxidative DNA damage. These functional changes were paralleled by distinct metabolomic profiles between control and knockout cells, as identified through multivariate analysis and pathway enrichment. Notably, ALDH2 deficiency significantly alters pathways involved in arginine biosynthesis and arginine, proline metabolism, glutathione metabolism, and nicotinate/nicotinamide metabolism. These findings verified that ALDH2 serves as a critical regulator of a stress-associated metabolic axis that links aldehyde detoxification capacity to nitrogen metabolism, redox homeostasis, and mitochondrial lipid utilization

ALDH2 deficiency is known to elevate aldehyde stress and oxidative injury across tissues and disease contexts [15]. Our functional characterization of isogenic SKBR-3 sublines shows that ALDH2 deletion increased the burden of endogenously generated reactive aldehydes from basal and stress-enhanced lipid peroxidation, promoting a feed-forward cycle of mitochondrial dysfunction and oxidative stress (Fig. 2), which is consistent with prior studies [14]. Since cells used in this study were not subjected to alcohol or other stimulants, our findings suggest that ALDH2 loss exerts significant cellular effects through endogenous lipid peroxidation-derived aldehydes, such as 4-HNE, in addition to acetaldehyde. This is supported by studies in Aldh2 Glu504Lys knock-in mice showing increased 4-HNE-adducted proteins enriched in mitochondrial fatty acid oxidation and electron transport pathways, accompanied by impaired mitochondrial respiration [21]. Although we did not directly quantify 4-HNE or aldehyde adducts, the observed increases in MDA and ROS are consistent with enhanced lipid peroxidation and reduced carbonyl detoxification capacity. Together, these data links ALDH2 deficiency to aldehyde adduct accumulation on mitochondrial proteins and disruption of mitochondrial energy metabolism, also highlighting the toxic effects of endogenous aldehydes.

Our metabolomics analyses suggest that a major adaptive program to ALDH2 loss involves reprogramming arginine/urea cycle metabolism toward polyamine-associated pathways. We observed accumulation of L-ornithine and L-arginine coupled to depletion of L-citrulline and argininosuccinic acid in ALDH2 knockout cells, supporting impaired flux through the argininosuccinate synthase/lyase segment of arginine biosynthesis/urea-cycle-linked reactions (Fig.6). In parallel, polyamine balance shifted toward short polyamines (increased putrescine) with depletion of spermine. This pattern is notable because urea cycle enzyme expression and metabolite flow are frequently dysregulated in cancer in a context-dependent manner, with consequences for biosynthesis, tumor growth, and tumor immune interactions [22]. Moreover, an explicit urea-cycle/polyamine axis has been implicated as a cancer-promoting metabolic configuration in other tumor settings [23]. Our data therefore support a model in which ALDH2 deficiency biases nitrogen handling away from “disposal/synthesis” routes that generate citrulline/argininosuccinate and toward polyamine-associated utilization of ornithine, potentially favoring proliferative and stress-adaptive phenotypes. Importantly, the connection between ALDH2 and arginine handling is increasingly recognized in the cancer metabolism literature. In colorectal cancer, metabolomic profiling has identified ALDH2 as a regulator of intracellular L-arginine turnover through an ALDH2/ARG2 axis, with therapeutic implications for immunotherapy responsiveness [24]. Although our breast cancer model shows a modest increase in arginine with decreased downstream urea-cycle metabolites, both datasets indicate that ALDH2 governs arginine pool balance and metabolic routing in cancer. One possible mechanism is that ALDH2-dependent aldehyde/redox stress influences the expression or activity of key arginine-pathway enzymes (e.g., ASS1/ASL, arginases, or polyamine synthesis/catabolism enzymes), thereby changing flux even when bulk arginine abundance rises. Direct enzyme/protein quantification, isotope tracing, and NO-related readouts would clarify whether altered arginine utilization contributes to oxidative stress amplification or represents an attempted compensatory response.

Our data also indicated that several carnitine derivates were downregulated in ALDH2 knockout cells, suggesting that reduced carnitine levels may contribute to the increased oxidative stress induced by ALDH2 deficiency. This coordinated reduction may represent a metabolic signature of impaired mitochondrial lipid utilization, indicating that ALDH2 deficiency disrupts fatty acid transport and mitochondrial function. Previous report has shown that some ALDH family members, such as ALDH9, ALDH25, ALDH26, and ALDH27, exhibit ω-aminoaldehyde dehydrogenases activity, involved in carnitine synthesis [25]. Mitochondrial ALDH2 has also been implicated in this pathway [26], although whether its deficiency directly impairs carnitine biosynthesis remains unclear. In addition, altered lipid peroxidation and mitochondrial stress may further disturb carnitine homeostasis, linking ALDH2 to lipid metabolic regulation. Carnitine is essential for transporting long-chain fatty acids into mitochondria for β-oxidation and thus plays a key role in energy metabolism [27]. Disruption of this process can promote lipid accumulation and oxidative stress. Moreover, carnitine has recognized antioxidant properties [28], and L-carnitine supplementation has been shown to mitigate oxidative damage [29,30]. Together, these findings support a role for carnitine as a lipid/mitochondrial signature in ALDH2 deficiency and warrant further investigation into its contribution to ALDH2-mediated antioxidant effects.

## 5. Conclusion

In summary, ALDH2 knockout in SKBR-3 cells induces oxidative stress and oxidative DNA damage and drives coordinated metabolic remodeling that prominently involves arginine/urea-cycle-linked metabolism, polyamine balance, glutathione pathways, and mitochondrial lipid-associated metabolites. These data support a model in which ALDH2 deficiency rewires cancer cell metabolism in response to aldehyde/redox stress, creating both pro-damage conditions and potentially targetable metabolic dependencies.

## Supporting information

Normalized Data with metadata

Differentialed metabolites

Metabolites with VIP more than 1

Enriched pathways

Metabolites in top 5 pathways

## Data availability

All data are included in the manuscript and its supplementary files.

